# Topology-dependent FRET efficiency in living cells via N-C swapping of fluorescent protein fusions

**DOI:** 10.64898/2026.06.03.729767

**Authors:** Takashi Tanida, Md. Royhan Gofur, Takayuki Nakajima

## Abstract

Förster resonance energy transfer (FRET) is a physicochemical phenomenon involving non-radiative energy transfer between donor and acceptor fluorophores. While FRET efficiency primarily depends on the proximity between fluorophores, additional factors substantially influence the efficiency in living cells. However, how non-distance factors modulate live-cell FRET efficiency remains poorly understood. Here, we report the significant role of N-and C-terminal topology in determining live-cell FRET efficiency, independent of fluorophore proximity, donor variants, and subcellular compartment. Using acceptor photobleaching and sensitized emission measurements in living cells, we found that FRET efficiencies of mCherry-EGFP or mCherry-EYFP (acceptor-donor) were significantly higher than those of EGFP-mCherry or EYFP-mCherry (donor-acceptor), respectively. FRET efficiencies were higher with EYFP than with EGFP as the donor. These efficiencies were comparable between the nucleus and cytoplasm. AlphaFold2-based structural modeling suggested similar proximity between donor and acceptor fluorophores despite structurally heterogeneous and loosely constrained geometries. In contrast, ensemble FRET simulations suggested that dynamic conformational sampling amplifies the apparent asymmetry between donor-acceptor and acceptor-donor configurations observed in live-cell FRET measurements. Collectively, these results demonstrate that topological arrangement, rather than static structural geometry alone, plays a significant role in FRET efficiency in living cells, providing molecular implications for the design of intramolecular FRET-based biosensors.

## Introduction

Förster resonance energy transfer (FRET) is a non-radiative energy transfer process between fluorophores mediated by dipole–dipole coupling [1–3]. Excitation energy from a fluorophore with a shorter emission wavelength, the donor, is transferred to a fluorophore with a longer emission wavelength, the acceptor, when the donor emission spectrum overlaps with the acceptor absorption spectrum [1–3]. FRET largely depends on the proximity between those two fluorophores, typically within 10–100 Å, and therefore it serves as a “molecular ruler” between two molecules [4–6]. Given that typical proteins are tens of nanometers in size, FRET can reveal direct protein–protein interactions in biochemical assays. FRET microscopy is widely applied for biophysical measurements of proteins-of-interest tagged with fluorescent proteins in living cells [7,8]. These FRET-based techniques enable the visualization of real-time subcellular interactions of proteins in living cells [7–11]. Various biosensors have been developed to monitor the dynamics of subcellular biomolecules and pathophysiological conditions in living cells. These biosensing tools rely on conformational changes in response to the subcellular environment or ligand stimulation, which induce intramolecular interaction and subsequent FRET-mediated spectral shift [12–15].

The efficiency of energy transfer from the donor to acceptor, which occurs alongside the donor fluorescence decay, is defined as the FRET efficiency. This index is inversely proportional to the sixth power of the distance between the donor and acceptor. In addition to proximity, the dipole orientation and spectral overlap are other critical determinants of FRET efficiency. The dipole orientation, referred to as the orientation factor (*κ*^2^), ranges from 0 to 4 and fundamentally governs the energy transfer rate in FRET [16,17]. This factor is determined by transition dipole moment orientations between FRET probe pairs. Typical macromolecules such as fluorescently labeled proteins are often assumed to rotate freely and isotropically in aqueous media, and thus a value of *κ*^2^ = 2/3 is conventionally used in many studies [18,19].

Another factor influencing FRET efficiency is the spectral overlap, *J*(λ), between donor emission and acceptor absorbance [18,20]. While a larger spectral overlap increases FRET efficiency, it simultaneously complicates the discrimination between donor and acceptor fluorescence because of significant bleed-through in detection systems [18,19]. Furthermore, subcellular compartments and organelles present diverse microenvironments, characterized by varying pH, metabolite concentrations, and macromolecular crowding, all of which can modulate FRET efficiency [20–23]. Consequently, FRET signals are determined not only by donor-acceptor distances but also by the interplay of orientation factors, spectral characteristics, and the local subcellular environment [5]. However, the quantitative measurement and validation of the influence of these factors on FRET efficiency in living cells remain insufficiently addressed.

Computational modeling has become a powerful tool for understanding the structural basis of protein function. AlphaFold enables high-accuracy predictions of 3D protein structures [24–27], potentially enabling the estimation of the relative spatial arrangement and orientation of fluorophores within FRET sensor fusion proteins. Integrating AlphaFold-based structural modeling with experimental FRET measurements [26,27] may offer a useful framework for interpreting how the topological arrangements of the probe influences energy transfer efficiency beyond simple linear distance.

The relative topology of donor and acceptor domains may contribute to distance-independent modulation of FRET efficiency. To investigate how non-proximity factors influence FRET efficiency in living cells, we used a simplified model system comprising N-and C-terminally (N–C) swapped donor–acceptor constructs linked via a conserved flexible linker, closely related donor variants, and differential subcellular localization (nucleus vs. cytoplasm). This approach enabled quantitative evaluation of non-distance effects under conditions where the intermolecular distance between fluorophores is effectively constrained. Our results reveal that FRET efficiency is consistently modulated by topological arrangements, independent of donor identity and subcellular localization. Furthermore, we used AlphaFold2-based structural predictions together with ensemble FRET simulations to interpret the experimental observations in the context of both static and dynamic structural models. Collectively, our findings provide fundamental molecular insights into the role of topological factors in energy transfer efficiency, with implications for the design of effective intramolecular FRET sensors in living cells.

## Results

### Acceptor photobleaching FRET microscopy in living cells

To investigate the factors that influence living cell FRET efficiency other than fluorophore proximity, we generated plasmid constructs with donor-acceptor (DA) configuration, EGFP-mCherry and EYFP-mCherry, and their counterparts with acceptor-donor (AD) configuration, mCherry-EGFP and mCherry-EYFP (Fig. 1). The donor and acceptor pairs were connected by an 18-amino-acid flexible linker and expressed under an identical CMV promoter derived from the pEYFP or pEGFP backbone.

**Figure 1.**
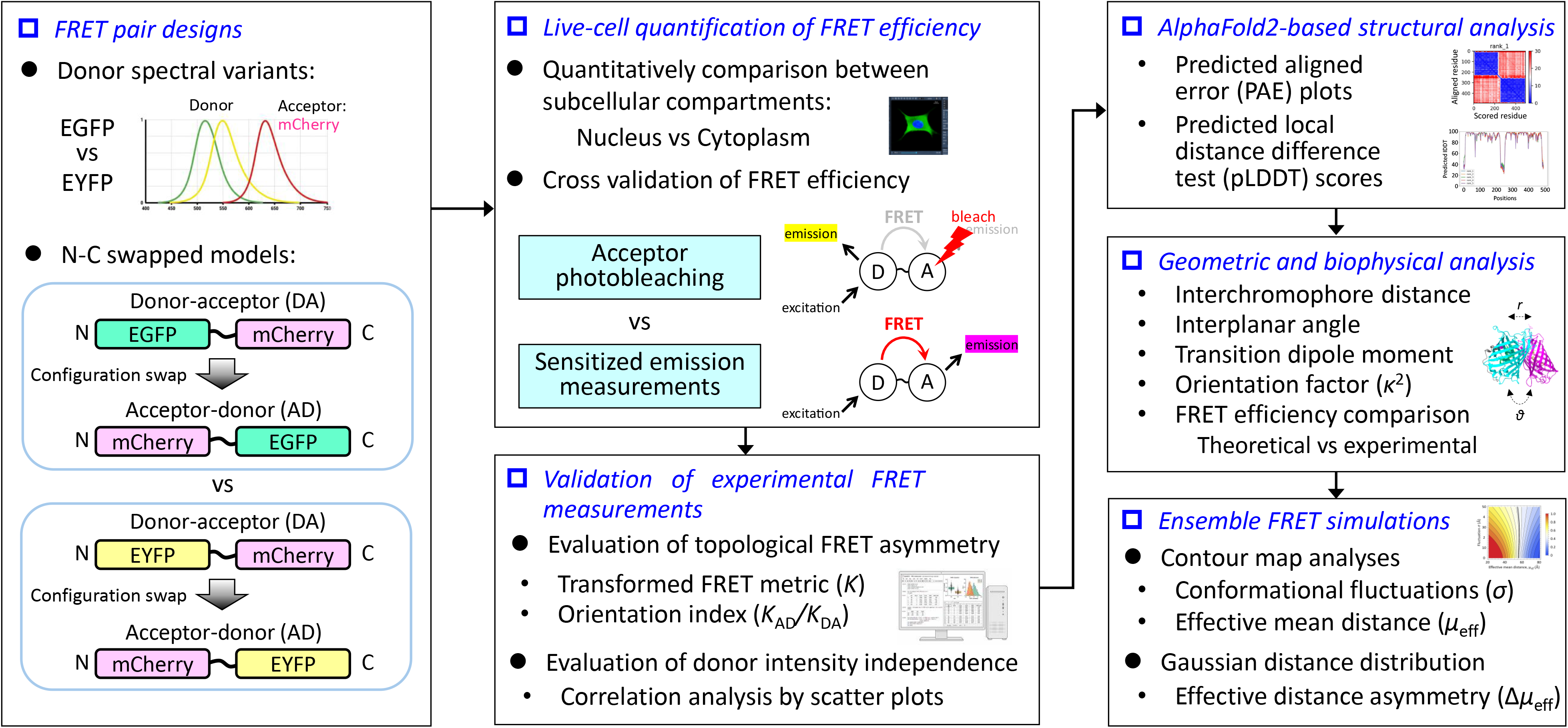
Workflow for validating topology-dependent FRET using N–C terminal swapped constructs. FRET efficiencies of N–C terminal swap models with donor spectral variants of EGFP and EYFP were analyzed by acceptor photobleaching and sensitized emission measurements separately in the nucleus and cytoplasm of living cells. The observed efficiencies were analyzed theoretically, and independence from donor fluorescence intensity was confirmed using correlation coefficients. *In silico* structural analysis was performed using 3D prediction models by AlphaFold. Theoretical values of inter-chromophore distance, angular geometry, transition dipole moment, and orientation factor (*κ*^2^) were calculated using ColabFold and UCSF ChimeraX, and theoretical FRET efficiencies were compared with experimental values. Ensemble FRET simulations were performed in Wolfram Mathematica using the experimentally observed FRET efficiencies together with the structure-based Förster distances (*R*_0_).

Acceptor photobleaching FRET microscopy is a widely accepted method to measure FRET efficiency (Fig. 1). This approach is based on the increase of donor fluorescence following photobleaching-induced loss of acceptor fluorescence (Fig. 1). With both DA and AD configurations, donor fluorescence intensity markedly increased after acceptor photobleaching in COS-1 cells, while the non-fused control (D/A) where fluorophores were expressed separately showed no detectable donor dequenching (Fig. 2a, b).

**Figure 2.**
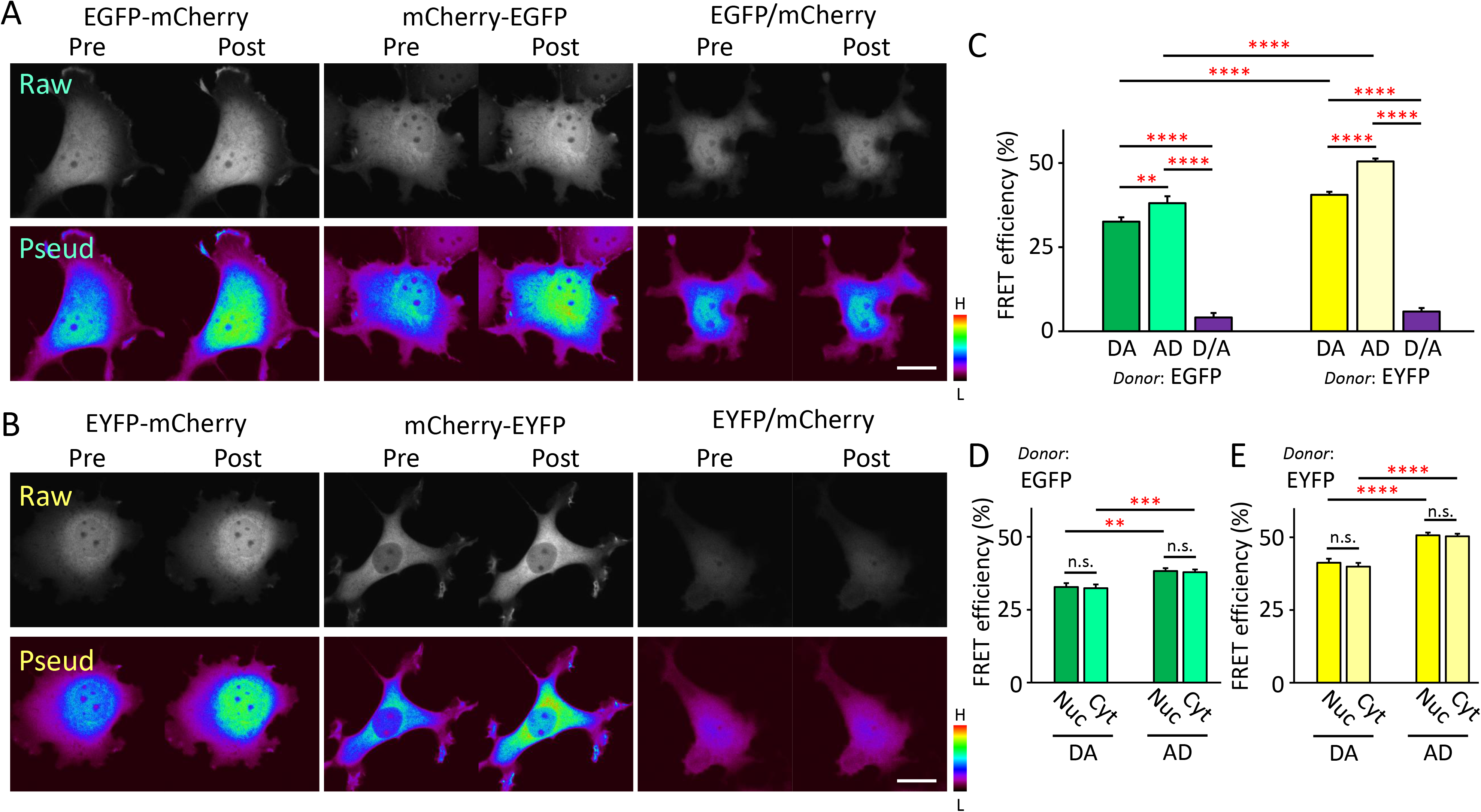
Live-cell acceptor photobleaching FRET microscopy. a, b. Raw (*top*) and pseudocolored (*bottom*) confocal images of donor fluorescent protein (a: EGFP or b: EYFP) in COS-1 cells expressing the indicated fluorophore combinations before (*Pre*) and after (*Post*) photobleaching of acceptor protein (mCherry). COS-1 cells were transiently transfected with EGFP-mCherry and mCherry-EGFP (a) or EYFP-mCherry and mCherry-EYFP (b) expression vectors. As negative controls, the EGFP (a) or EYFP (b) expression vector was co-transfected with an expression vector encoding non-fused mCherry. After transfection, live-cell imaging was performed using a confocal laser scanning microscope. Experiments were repeated at least three times. Pseudocolor bars indicate fluorescence intensity, with H and L representing high and low signal intensity, respectively. Bar = 20 µm. c. FRET efficiency determined by acceptor photobleaching in COS-1 cells expressing EGFP-mCherry (n = 31), mCherry-EGFP (n = 34), EYFP-mCherry (n = 28), or mCherry-EYFP (n = 33). As negative controls, COS-1 cells expressing non-fused combinations of EGFP and mCherry (n = 28) or EYFP and mCherry (n = 27) were analyzed. Data were obtained from three independent experiments. d, e. FRET efficiencies determined by live-cell acceptor photobleaching in the nucleus or cytoplasm of COS-1 cells expressing EGFP-mCherry (n = 31 each for nucleus and cytoplasm) and mCherry-EGFP (n = 34 each for nucleus and cytoplasm each) (d), or EYFP-mCherry (n = 28 each for nucleus and cytoplasm) and mCherry-EYFP (n = 33 each for nucleus and cytoplasm) (e). Data were obtained from three independent experiments. *DA*, donor-acceptor configuration (EGFP-mCherry and EYFP-mCherry); *AD*, acceptor-donor configuration (mCherry-EGFP and mCherry-EYFP); *D/A*, non-fused control of donor (EGFP or EYFP) and acceptor (mCherry); *Nuc*, nucleus; *Cyt*, cytoplasm. Values are expressed as means ± SE. \**p* < 0.05, \*\**p* < 0.01, \*\*\**p* < 0.001, \*\*\*\**p* < 0.0001 by two-way ANOVA followed by Bonferroni/Dunn’s post-*hoc* tests. n.s., not significant.

Fluorescence intensities were measured from regions of interest (ROIs) placed separately in the nucleus and cytoplasm. For accurate quantification of FRET efficiency in the acceptor photobleaching, fluorescence intensities and the efficiencies were corrected using background intensities in each image and bleaching efficiency, respectively (Equation 1). Using mCherry-expressing cells, the averaged bleaching efficiencies were 69.2 ± 0.5% in the nucleus and 67.5 ± 0.6% in the cytoplasm (Supplementary Fig. S1).

Two-way ANOVA analysis indicated that both configuration (DA, AD, or D/A) and donor (EGFP vs. EYFP) significantly affected the FRET efficiency (configuration: *F*(2, 175) = 524.30, *p* < 0.0001; donor: *F*(1, 175) = 50.80, *p* < 0.0001). The interaction between donor and configuration was also significant, suggesting the difference of sensitivity to N–C configuration between EGFP and EYFP as donor (*F*(2, 175) = 8.85, *p* = 0.0002).

To further characterize the differences between configurations and donors, post-*hoc* multiple comparisons were performed. In the comparison of donor spectral variants, significantly higher FRET efficiencies were observed for EYFP-mCherry and mCherry-EYFP compared with their configuration-matched constructs, EGFP-mCherry and mCherry-EGFP, respectively (EYFP-mCherry vs. EGFP-mCherry: *p* < 0.0001; mCherry-EGFP vs. mCherry-EYFP, *p* < 0.0001) (Fig. 2c). The FRET efficiencies for mCherry-EGFP (38.1 ± 0.9%) and mCherry-EYFP (50.5 ± 0.8%) were significantly higher than those for EGFP-mCherry (32.6 ± 1.3 %) and EYFP-mCherry (40.6 ± 2.1%), respectively (constructs with EGFP donor: *p* = 0.0017; constructs with EYFP donor: *p* < 0.0001), indicating that the AD configuration (mCherry-EGFP and mCherry-EYFP) yielded higher FRET efficiency compared with the DA configuration (EGFP-mCherry and EYFP-mCherry) (*p* < 0.0001) (Fig. 2c). In the non-fused control constructs, the FRET efficiencies were hardly detected (EGFP/mCherry: 4.1 ± 1.3%; EYFP/mCherry: 5.9 ± 1.0%), indicating the validity of this experimental system of FRET microscopy (Fig. 2c).

Subcellular locations including nucleus, cytoplasm, and organelles have various unique microenvironments arising from differences in pH, molecular crowding, metabolites, and electrolytes. We next investigated whether FRET efficiency is affected by the subcellular location (nucleus and cytoplasm) in living cells.

For the EGFP constructs, two-way ANOVA revealed that the subcellular compartment (nucleus and cytoplasm) had no statistical effects (*F*(1, 126) = 0.112, *p* = 0.738), although the configuration effect was markedly significant (*F*(1, 126) = 23.63, *p* < 0.0001). An interaction between configuration and compartment was also not detected (*F*(1, 126) = 2.91 × 10^-4^, *p* = 0.986). Further multiple comparison tests revealed that the mCherry-EGFP (nucleus: 38.3 ± 1.0%; cytoplasm: 37.9 ± 0.9%) yielded significantly higher FRET efficiency than EGFP-mCherry (nucleus: 32.8 ± 1.4%; cytoplasm: 32.4 ± 1.3%) in both the nucleus and cytoplasm when each compartment was analyzed separately (nucleus: *p* = 0.0013; cytoplasm: *p* = 0.0008) (Fig. 2d).

Consistent findings were observed for the EYFP constructs. The subcellular compartment (nucleus vs. cytoplasm) and the interaction between configuration and compartment elicited no statistically significant effects (compartment: *F*(1, 118) = 0.329, *p* = 0.568; interaction: *F*(1, 118) = 0.103; *p* = 0.749). However, we observed highly significant effects in the configuration (EYFP-mCherry vs. mCherry-EYFP, *F*(1, 118) = 43.28, *p* < 0.0001). Further multiple comparisons demonstrate that mCherry-EYFP (nucleus: 50.7 ± 0.8%; cytoplasm: 50.3 ± 0.9%) had significantly higher FRET efficiency than EYFP-mCherry (nucleus: 41.2 ± 2.2%; cytoplasm: 39.9 ± 2.0%) in both the nucleus (*p* < 0.0001) and cytoplasm (*p* < 0.0001) when each compartment was analyzed separately (Fig. 2e).

Thus, the superiority in FRET efficiency of the AD configuration compared with the DA configuration was observed. It was conserved in donor spectral variants, EGFP and EYFP, and not affected by subcellular environment, between the nucleus and cytoplasm. Furthermore, constructs containing EYFP as the donor exhibited significantly higher FRET efficiencies than the corresponding EGFP-containing constructs across all configurations and subcellular locations, consistent with the greater spectral overlap between EYFP and mCherry compared with EGFP and mCherry.

### Sensitized emission measurements of FRET efficiency in living cells

To cross-validate the FRET efficiency values obtained by acceptor photobleaching, we performed sensitized emission analysis based on the acceptor-to-donor fluorescence intensity ratio. A clear increase in acceptor fluorescence was observed in all fused constructs (EGFP-mCherry, EYFP-mCherry, and their corresponding N–C swapped variants) compared with non-fused controls (Fig. 3a, b). For accurate quantification of FRET efficiency, fluorescence intensities were corrected for background and spectral bleed-through (Equation 2). Donor bleed through of EGFP and EYFP was corrected while acceptor bleed through of mCherry was not detected in the system and accordingly it was not incorporated to corrections for FRET efficiency. The γ factor, determined by acceptor photobleaching, was applied to correct for differences in detection efficiency and quantum yield between donor and acceptor (Equations 3, 4).

**Figure 3.**
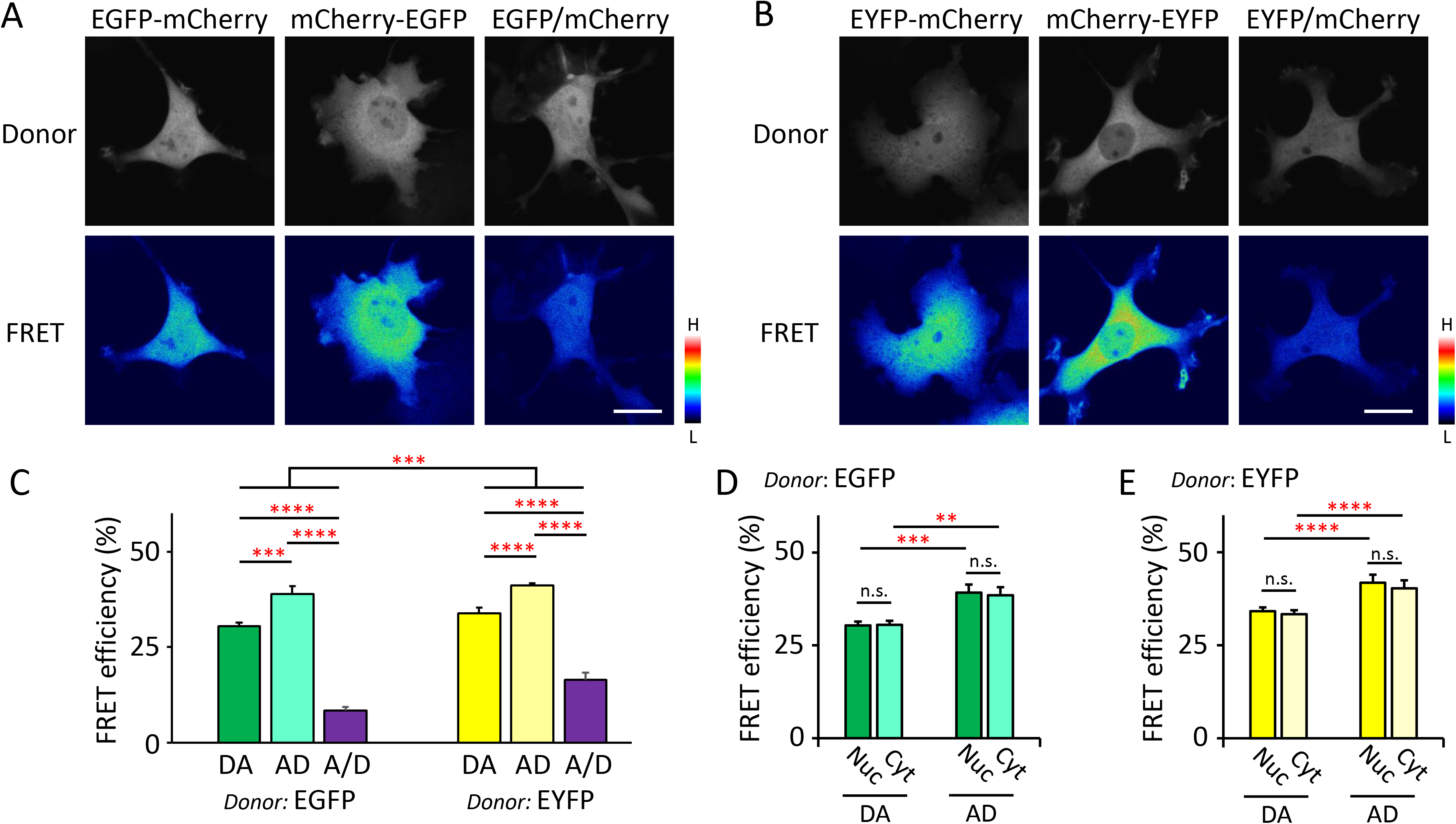
Sensitized emission FRET measurements in living cells a, b. Confocal live-cell images of donor fluorescent protein (a: EGFP or b: EYFP) and FRET signal channels in COS-1 cells expressing the indicated fluorophore combinations. COS-1 cells were transiently transfected with EGFP-mCherry and mCherry-EGFP (a) or EYFP-mCherry and mCherry-EYFP (b) expression vectors. As negative controls, the EGFP (a) or EYFP (b) expression vector was co-transfected with an expression vector encoding non-fused mCherry. After transfection, live-cell imaging was performed using a confocal laser scanning microscope. Experiments were repeated at least three times. Pseudocolor bars indicate fluorescence intensity, with H and L representing high and low signal intensity, respectively. Bar = 20 µm. c. FRET efficiency determined by sensitized emission in living COS-1 cells expressing EGFP-mCherry (n = 31), mCherry-EGFP (n = 34), EYFP-mCherry (n = 27), or mCherry-EYFP (n = 33). COS-1 cells expressing non-fused combinations of EGFP and mCherry (n = 25) or EYFP and mCherry (n = 22) served as negative controls. Data were obtained from three independent experiments. d, e. FRET efficiencies determined by live-cell sensitized emission in the nucleus (*Nuc*) or cytoplasm (*Cyt*) of COS-1 cells expressing EGFP-mCherry (n = 31 each for nucleus and cytoplasm) and mCherry-EGFP (n = 34 each for nucleus and cytoplasm) (d), or EYFP-mCherry (n = 27 each for nucleus and cytoplasm) and mCherry-EYFP (n = 33 each for nucleus and cytoplasm) (e). Data were obtained from three independent experiments. *DA*, donor-acceptor configuration (EGFP-mCherry and EYFP-mCherry); *AD*, acceptor-donor configuration (mCherry-EGFP and mCherry-EYFP); *D/A*, non-fused control of donor (EGFP or EYFP) and acceptor (mCherry); *Nuc*, nucleus; *Cyt*, cytoplasm. Values are expressed as means ± SE. \*\**p* < 0.01, \*\*\**p* < 0.001, \*\*\*\**p* < 0.0001 by two-way ANOVA followed by Bonferroni/Dunn’s post-*hoc* tests. n.s., not significant.

Consistent with acceptor photobleaching (Fig. 2), two-way ANOVA analysis revealed markedly significant effects in configuration (DA vs. AD) and donor (EGFP vs. EYFP) (configuration: *F*(2, 166) = 171.49, *p* < 0.0001; donor: *F*(1, 166) = 13.61, *p* = 0.0003). However, an interaction between configuration and donor was not detected (*F*(2, 166) = 1.91, *p* = 0.151). Further multiple comparison tests demonstrated that mCherry-EGFP (38.9 ± 2.1%) and mCherry-EYFP (41.1 ± 0.7%) yielded significantly larger FRET efficiency compared with their corresponding N–C swaps, EGFP-mCherry (30.5 ± 1.1%) and EYFP-mCherry (33.8 ± 1.5%), respectively (EGFP constructs: *p* = 0.0002; EYFP constructs: *p* < 0.0001) (Fig. 3c). Thus, sensitized emission analysis revealed that the AD configuration effectively generates FRET efficiency compared with the DA configuration, consistent with acceptor photobleaching (Fig. 2).

To cross-validate the difference of FRET efficiency within the cellular compartment shown by acceptor photobleaching, we separately calculated FRET efficiency within the nucleus and cytoplasm in this sensitized emission method. In the two-way ANOVA analysis within the EGFP constructs, configuration (EGFP-mCherry vs. mCherry-EGFP) exhibited a significant effect (*F*(1, 126) = 23.04, *p* < 0.0001). However, compartment and interaction between configuration and compartment displayed no significant effect (compartment: *F*(1, 126) = 0.0267, *p* = 0.871; interaction: *F*(1, 126) = 0.0637, *p* = 0.801). Further multiple comparison tests revealed that mCherry-EGFP showed significantly larger FRET efficiency than EGFP-mCherry within both the nucleus (EGFP-mCherry: 30.4 ± 1.1% and mCherry-EGFP: 39.2 ± 2.1%; *p* = 0.0006) and cytoplasm (EGFP-mCherry: 30.5 ± 1.1% and mCherry-EGFP: 38.5 ± 2.2%; *p* = 0.0023) (Fig. 3d). However, FRET efficiency was comparable between the nucleus and cytoplasm for both EGFP-mCherry (*p* = 0.919) and mCherry-EGFP (*p* = 0.812) (Fig. 3d).

Similar results were obtained for the EYFP constructs. In the two-way ANOVA analysis, configuration (EYFP-mCherry vs. mCherry-EYFP) had a significant effect (*F*(1, 116) = 44.12, *p* < 0.0001), while compartment and interaction between configuration and compartment exhibited no significant effect (compartment: *F*(1, 116) = 1.11, *p* = 0.294; interaction: *F*(1, 116) = 0.126, *p* = 0.724). Further multiple comparison tests revealed that mCherry-EYFP exhibited significantly larger FRET efficiency than EYFP-mCherry in both the nucleus (EYFP-mCherry: 34.2 ± 1.6% and mCherry-EYFP: 41.8 ± 0.7%; *p* < 0.0001) and cytoplasm (EYFP-mCherry: 33.4 ± 1.4% and mCherry-EYFP: 40.3 ± 0.7%; *p* < 0.0001) (Fig. 3e). However, FRET efficiency was comparable between the nucleus and cytoplasm for both EYFP-mCherry (*p* = 0.719) and mCherry-EYFP (*p* = 0.112) (Fig. 3e).

These sensitized emission measurements demonstrated similar results to the acceptor photobleaching method in the FRET efficiency (Fig. 2), indicating the superiority of the AD configuration compared with the DA configuration that is conserved in donor variants (EGFP and EYFP) and subcellular compartments (nucleus and cytoplasm).

### Evaluation of transformed FRET metric and orientation index

To facilitate comparison between configurations, we transformed FRET efficiency *E* into *K* = *E*/(1 − *E*), a monotonic transformation that expands the dynamic range [3]. Hereafter, *K* is referred to as the transformed (*t*) FRET metric. Similar transformations have been used to linearize FRET efficiency and enhance sensitivity to differences [28,29]. From this transformation, we defined an orientation index (*OI*) as *OI* = *K*_AD_/*K*_DA_, to quantify topology-dependent asymmetry between configurations.

The *t*FRET metric of AD configurations were significantly larger than those of DA configurations in any experimental conditions of both donor (EGFP, EYFP) and subcellular compartment (nucleus, cytoplasm), indicating the higher FRET efficiencies for the AD configuration than for the DA configurations (EGFP donor, nucleus: EGFP-mCherry, 0.472 ± 0.019; mCherry-EGFP, 0.614 ± 0.020; *t*(61) =-5.24, *p* < 0.0001; cytoplasm: EGFP-mCherry, 0.470 ± 0.022; mCherry-EGFP, 0.603 ± 0.018; *t*(61) =-4.65, *p* < 0.0001; EYFP donor, nucleus: EYFP-mCherry, 0.749 ± 0.048; mCherry-EYFP, 1.05 ± 0.033; *t*(59) =-5.20, *p* < 0.0001; cytoplasm: EYFP-mCherry, 0.702 ± 0.043; mCherry-EYFP, 1.03 ± 0.034; *t*(59) =-6.13, *p* < 0.0001) (Fig. 4a–d).

**Figure 4.**
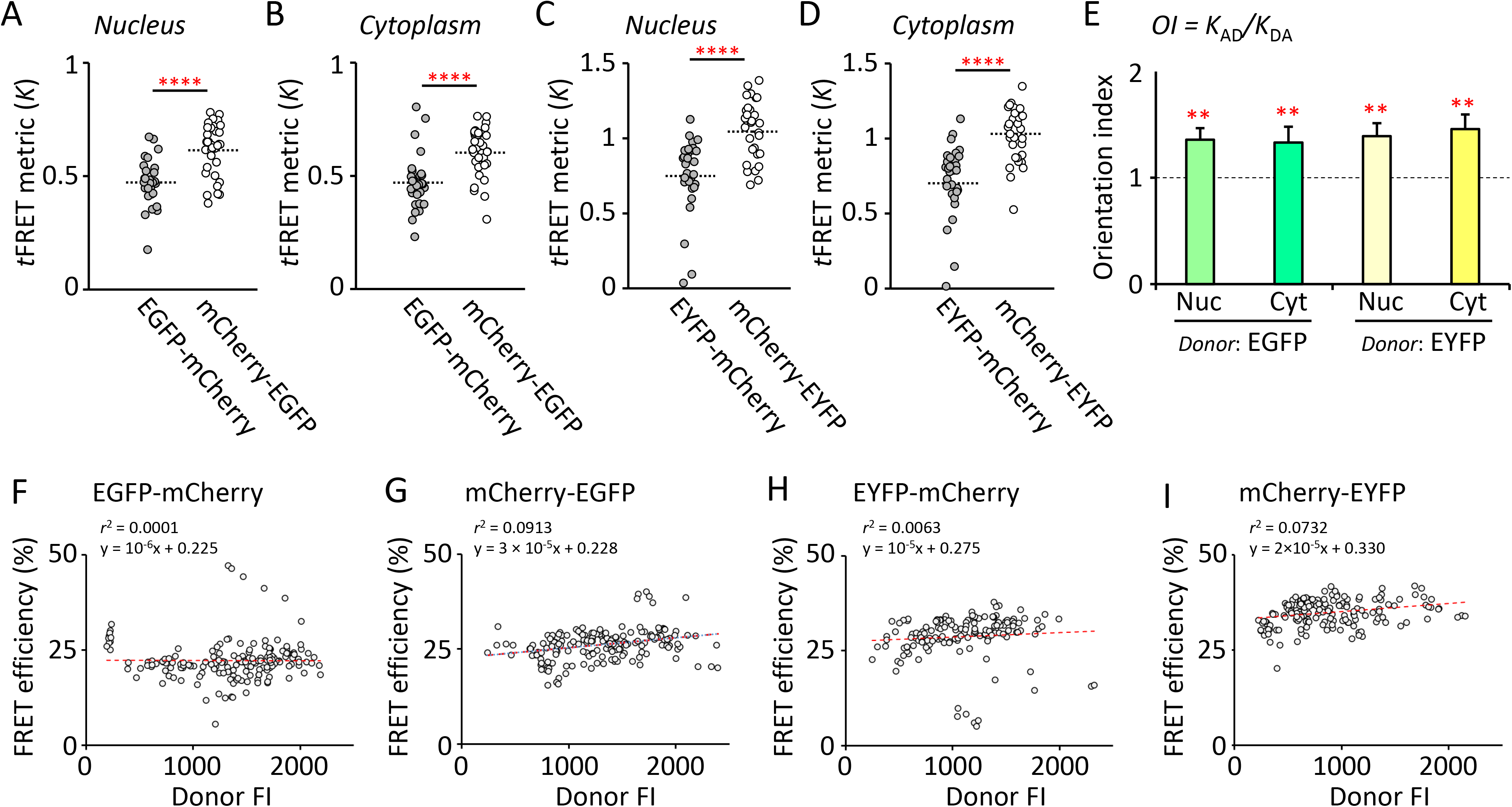
Evaluation of transformed FRET metric, orientation index, and independence from donor fluorescence intensity a–d. Jittered plots of the transformed (*t*) FRET metric, *K* = *E*/(1-*E*), calculated from FRET efficiency measured by live-cell acceptor photobleaching in COS-1 cells expressing the indicated fusion proteins: EGFP-mCherry (n = 30 each for nucleus and cytoplasm), mCherry-EGFP (n = 33 each for nucleus and cytoplasm), EYFP-mCherry (n = 28 each for nucleus and cytoplasm), and mCherry-EYFP (n = 33 each for nucleus and cytoplasm). Values are derived from three independent experiments. Data are shown as individual points (jittered). The dotted lines indicate the mean. \*\*\*\**p* < 0.0001 by unpaired *t*-test. e. Orientation index (*OI* = *K*_AD_/*K*_DA_) in EGFP-mCherry and mCherry-EGFP, and EYFP-mCherry and mCherry-EYFP, calculated separately in the nucleus (*Nuc*) and cytoplasm (*Cyt*). Statistics were performed on mean values from three independent experiments (n = 3). Data are shown as means ± SE. \*\**p* < 0.01 by one-sample *t*-test (µ = 1). The dotted line indicates 1.0. f–i. Scatter plots of donor fluorescence intensity (*FI*) vs. FRET efficiency measured by acceptor photobleaching in COS-1 cells expressing EGFP-mCherry (F; n = 173 ROIs), mCherry-EGFP (G; n = 181 ROIs), EYFP-mCherry (H; n = 151 ROIs), and mCherry-EYFP (I; n = 171 ROIs). Red dotted lines indicate linear regression fits (*r*² values shown). Data are from three independent experiments. Negligible correlations were observed.

To provide a quantitative index of configurational asymmetry, *OI*, the ratio of *K*_AD_ to *K*_DA_, was calculated for each donor and compartment using averaged *K* values from three independent experiments. All *OI* values (*K*_AD_/*K*_DA_) were significantly greater than unity in a one-sample *t*-test against a theoretical mean of 1.0, indicating that *K*_AD_ was consistently higher than *K_DA_*. These findings were conserved in all experimental conditions for both donor variants (EGFP or EYFP) and subcellular compartments (nucleus or cytoplasm), suggesting that AD configurations provide higher apparent FRET efficiencies compared with DA configurations (EGFP donor: nucleus, *OI* = 1.36 ± 0.11, *t*(2) = 12.55, *p* = 0.0063; cytoplasm, *OI* = 1.34 ± 0.13, *t*(2) = 10.50, *p* = 0.0089; EYFP donor: nucleus, *OI* = 1.39 ± 0.12, *t*(2) = 11.43, *p* = 0.0076; cytoplasm, *OI* = 1.46 ± 0.14, *t*(2) = 10.54, *p* = 0.0089) (Fig. 4e).

### Independence of donor fluorescence intensity in the FRET efficiency

To confirm whether these FRET efficiencies were affected by donor protein expression level, we analyzed the correlation between donor fluorescence intensities and FRET efficiencies within each ROI in the cells. As shown in Fig. 4e–i, linear regression analysis revealed negligible relationships for all fusion proteins, indicating that the observed differences in the FRET efficiencies are not attributable to variations in donor expression levels or fluorescence intensities in living cells (EGFP-mCherry: *r*² = 0.0001, y = 1.0 × 10⁻⁶x + 0.225; mCherry-EGFP: *r*² = 0.0913, y = 3.0 × 10⁻⁵x + 0.228; EYFP-mCherry: *r*² = 0.0063, y = 1.0 × 10⁻⁵x + 0.275; mCherry-EYFP: *r*² = 0.0732, y = 2.0 × 10⁻⁵x + 0.330).

### Artificial intelligence–based prediction of structures using Alphafold2

Mathematical modeling obtained from 3D structure prediction is an important determinant of the biomedical functions of proteins-of-interest [30,31]. Conserved polar bias of AD configuration in FRET efficiency was observed in the live-cell experiments, and the 3D structure-based theoretical prediction was performed using AlphaFold2 [24] (Supplementary Fig. S2).

The predicted aligned error (PAE) maps generated by AlphaFold2 indicate that individual fluorescent protein domains (EGFP, EYFP, and mCherry) are structurally stable, as shown by the blue squares representing low PAE values within each domain (Fig. 5a–d and Supplementary Fig. S3). In contrast, PAE values around residue 250, between the donor (EGFP or EYFP) and acceptor (mCherry) domains, were generally elevated (white to red) reaching up to ∼30 Å in part, with moderately heterogeneous patterns (Fig. 5a–d and Supplementary Fig. S3). The predicted local distance difference test (pLDDT) map also indicated that the linker region is not static but samples multiple conformations (Fig. 5a–d). Therefore, these 3D structure analyses suggest the absence of a single well-defined interface and the presence of loosely constrained relative orientations within all the fluorescent fusion proteins.

**Figure 5.**
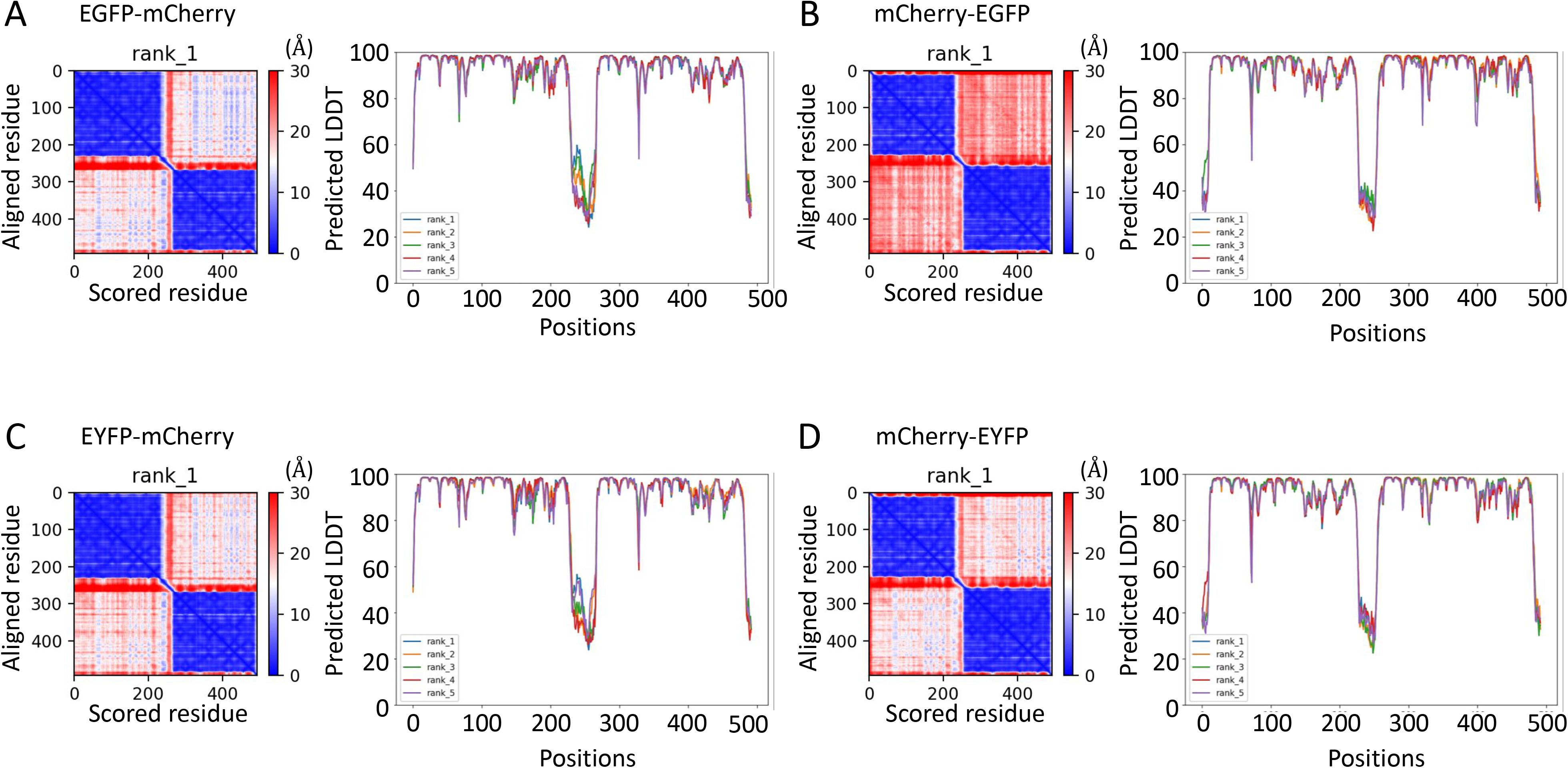
Structural characterization of predicted FRET pair models. a–d. Predicted aligned error (PAE) plots (*left*) of the top-ranked models and predicted local distance difference test (pLDDT) score profiles of the top five models (*right*) for EGFP-mCherry (a), mCherry-EGFP (b), EYFP-mCherry (c), and mCherry-EYFP (d), generated using ColabFold. Lower PAE values indicate higher confidence in the predicted relative positions of residue pairs, whereas higher pLDDT scores indicate greater confidence in local structural accuracy.

### Theoretical analysis using predicted 3D structure models

To evaluate the structural determinants of FRET efficiency, theoretical parameters were calculated from predicted 3D models generated by ColabFold [32] and analyzed using UCSF ChimeraX [33] (Fig. 6a–d). The donor-acceptor distances were highly conserved across all ranked models (constructs with EGFP as donor: 22.4–24.6 Å; constructs with EYFP as donor: 22.3–23.9 Å) with no significant differences (*F*(3, 16) = 1.62, *p* = 0.224; Fig. 6e and Supplementary Fig. S4a, b). Despite the flexible relative domain arrangement suggested by the PAE, the donor-acceptor distances remained narrowly distributed across models (*ΔR* ≈ 0.8–2.1 Å). Given the strong distance dependence of FRET efficiency [1,2], such small variations are unlikely to account for the observed differences in FRET. Thus, these similar donor-acceptor distances and poorly constrained relative fluorophore orientations inferred from AlphaFold2 (Fig. 5a–d) provide a structural framework suggesting that N–C topology contributes to the modulation of FRET efficiency in living cells beyond simple distance-dependent effects.

**Figure 6.**
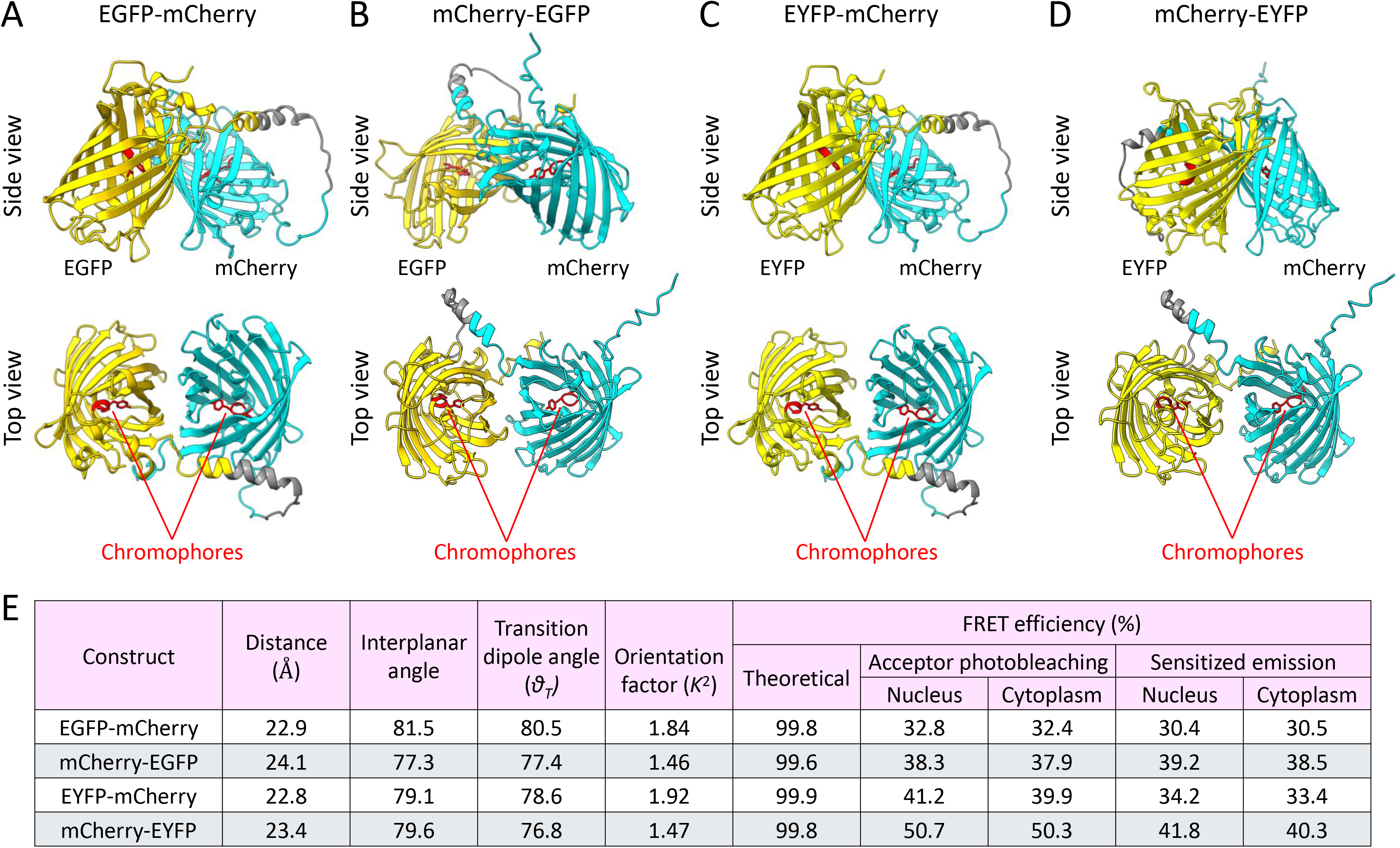
Predicted 3D models with theoretical structural metrics and FRET efficiencies. a–d. Side and top views of predicted 3D structures of EGFP-mCherry (a), mCherry-EGFP (b), EYFP-mCherry (c), and mCherry-EYFP (d) generated by ColabFold. Chromophores are highlighted in red. e. Summary of chromophore distances (Å), interplanar angles, transition dipole angles (*θ_T_*), orientation factors (*κ*^2^), and FRET efficiencies derived from AlphaFold2 predictions via ColabFold and structure analysis in ChimeraX, together with live-cell measurements from acceptor photobleaching and sensitized emission experiments.

We next analyzed non-distance factors in the top-ranked predicted structures using ChimeraX. Plane angles between the chromophores in these models were comparable across the four constructs (77.3– 81.5°; Fig. 6e). In contrast, the orientation factor (*κ^2^*) estimated from these structures using Equation 5 differed between the DA and AD configurations (EGFP-mCherry vs. mCherry-EGFP: Δ*κ^2^* = 0.38; EYFP-mCherry vs. mCherry-EYFP: Δ*κ^2^* = 0.45) (Fig. 6e). However, evaluations based on these top-ranked structures, incorporating both donor-acceptor distances and estimated orientation factors (*κ^2^*) (Equations 6, 7), yielded nearly saturated theoretical FRET efficiencies that differed markedly from experimental values (Figs. 2, 3 and 6e). When *κ^2^* varied from 0 to 4, the theoretical FRET efficiencies for all constructs remained close to saturation (> 99%) (Supplementary Fig. S4c). These discrepancies suggest that structural parameters alone at these proximities are insufficient to explain the observed live-cell FRET efficiencies, particularly the significantly higher FRET efficiency in the AD configuration compared with the DA configuration.

### Ensemble FRET simulations of conformational distance distributions

To investigate how conformational fluctuations and chromophore distance distributions contribute to the asymmetric FRET efficiencies observed between the DA and AD configurations, we generated pseudocolor contour maps describing the relationship between conformational fluctuation (*σ*), effective mean chromophore distance (*μ*_eff_), and theoretical ensemble FRET efficiency using Wolfram Mathematica (Fig. 7a–d; Supplementary Code S1) [34]. The structure-based Förster distances (*R*_0_) calculated using ColabFold and ChimeraX (Fig. 6e) together with the experimentally observed FRET efficiencies determined by acceptor photobleaching (Fig. 2) were used to generate the contour plots (Equation 8–10). These analyses enabled theoretical modeling of FRET efficiency under different degrees of conformational sampling.

**Figure 7.**
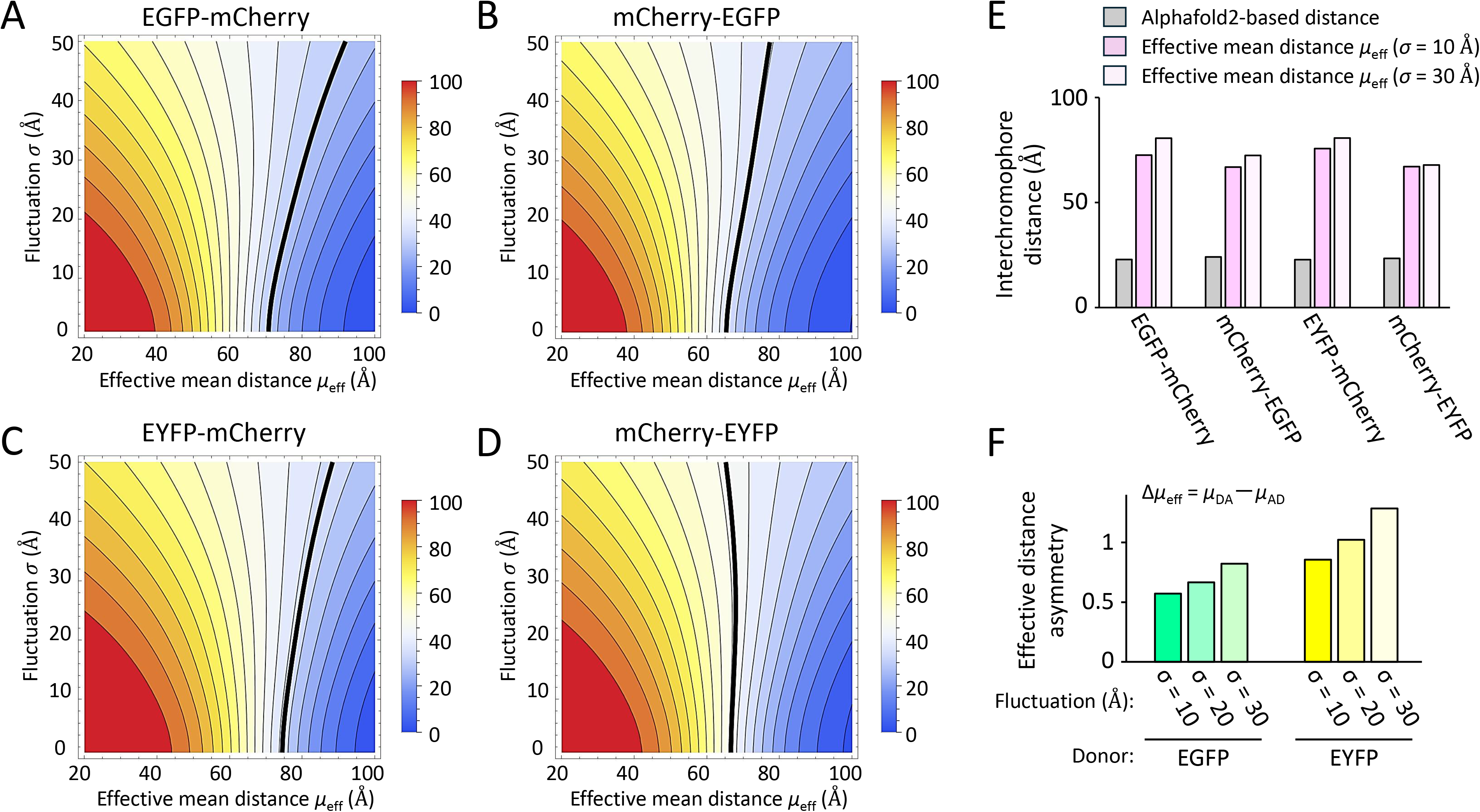
Ensemble FRET contour analysis for determining effective mean chromophore distances reproducing experimental FRET efficiencies. a–d. Pseudocolor contour plots showing theoretical ensemble FRET efficiencies as functions of effective mean chromophore distance (*μ*_eff_) and conformational fluctuation (*σ*) of the donor and acceptor domains for EGFP-mCherry (a), mCherry-EGFP (b), EYFP-mCherry (c), and mCherry-EYFP (d). The pseudocolor scales represent FRET efficiencies (0–100%). Black contour lines indicate the experimentally observed FRET efficiencies determined by acceptor photobleaching. e. Effective mean chromophore distances (*μ*_eff_) of EGFP-mCherry, mCherry-EGFP, EYFP-mCherry, and mCherry-EYFP at representative conformational fluctuations (*σ* = 10 Å and *σ* = 30 Å). Alphafold2-based chromophore distances are provided for comparison. f. Effective distance asymmetry (Δ*µ*_eff_) for EGFP and EYFP donor pairs at representative conformational fluctuations (*σ* = 10, 20, and 30 Å). Δ*μ*_eff_ was calculated as *μ*_DA_ − *μ*_AD_.

Regions with relatively small conformational fluctuations (*σ* < 20 Å) and short effective mean chromophore distances (*μ*_eff_ < 40 Å) corresponded to nearly saturated FRET efficiencies in all four constructs (Fig. 7a–d). The contour lines corresponding to the experimentally observed FRET efficiencies differed among the four constructs (Fig. 7a–d). When the effective mean chromophore distances (*μ*_eff_) required to reproduce the experimentally observed FRET efficiencies were calculated for relatively small (*σ* = 10 Å) and large (*σ* = 30 Å) conformational fluctuations, they were markedly larger than the AlphaFold-based static chromophore distances (Fig. 7e), suggesting that greater conformational fluctuations required larger *μ*_eff_ values to reproduce the experimentally observed FRET efficiencies. Additionally, the effective mean distances (*μ*_eff_) in the DA configuration were consistently larger than those in the AD configurations for a given *σ* (Fig. 7e). Therefore, the difference in effective mean chromophore distances (Δ*μ*_eff_ = *μ*_DA_ − *μ*_AD_) (Equation 11) was calculated for each donor fluorophore. Notably, Δ*μ*_eff_ remained positive in both EGFP and EYFP donor constructs, indicating that the effective mean chromophore distance was consistently greater in the DA configuration than in the AD configuration (Fig. 7f). Moreover, Δ*μ*_eff_ exhibited a progressive increase across the three *σ* values (10, 20, and 30 Å) for both EGFP-and EYFP-based donor pairs (Fig. 7f). This trend suggests that greater conformational fluctuations are associated with a larger DA/AD asymmetry in effective mean chromophore distance required to reproduce the live-cell FRET efficiencies.

### Analysis of effective mean distance asymmetry under conformational fluctuation

To examine whether the asymmetric relationship between conformational fluctuation and effective mean chromophore distance is a general property, we performed ensemble FRET simulations using Gaussian distance distributions generated in Mathematica for EGFP and EYFP donor constructs (Fig. 8a). As the assumed conformational fluctuation (*σ*) increased, the effective distance asymmetry (Δ*μ*_eff_ = *μ*_DA_ − *μ*_AD_) required to reproduce the experimentally observed FRET efficiencies increased progressively for both EGFP-and EYFP-based donor pairs (Fig. 8a). The Δ*μ*_eff_ values remained positive throughout the analyzed range and were consistently larger for EYFP donor pairs than for EGFP donor pairs (Fig. 8a), indicating a greater degree of DA/AD asymmetry in EYFP-based FRET pairs than in EGFP-based pairs. Based on these observations, larger conformational fluctuations are associated with progressively greater effective DA/AD distance asymmetry to reproduce the experimentally observed FRET efficiencies (Fig. 8b). Collectively, these results suggest that the apparent DA/AD asymmetry of the live-cell FRET efficiencies is determined predominantly by the extent of conformational sampling rather than by the absolute chromophore distance itself.

**Figure 8.**
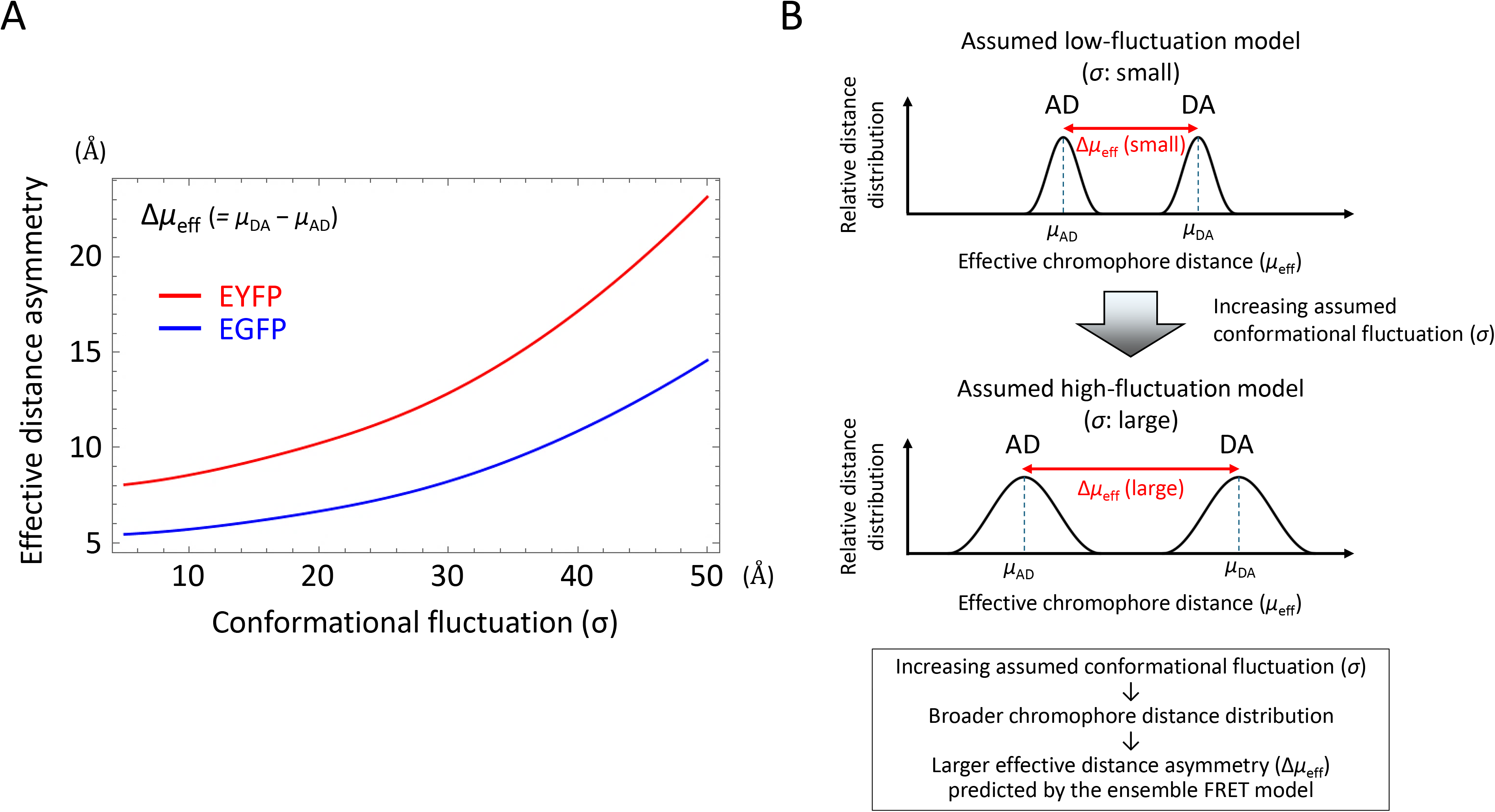
Effect of conformational fluctuation on effective DA/AD chromophore distance asymmetry in ensemble FRET simulation. a. Effective distance asymmetry (Δ*μ*_eff_ = *μ*_DA_ − *μ*_AD_) plotted as a function of conformational fluctuation (*σ*) for EGFP and EYFP donor pairs. Increasing conformational fluctuation progressively increased Δ*μ*_eff_ while maintaining positive values throughout the analyzed range. b. Schematic illustration of the relative chromophore distance distributions and effective mean chromophore distances for the DA and AD configurations under relatively small (*top*) and large (*bottom*) conformational fluctuations (*σ*). As the assumed conformational fluctuation (*σ*) increases, the chromophore distance distributions broaden, resulting in greater effective mean chromophore distance asymmetry (Δ*μ*_eff_) predicted by the ensemble FRET model.

## Discussion

In this study, we investigated the effect of N–C terminal topology on FRET efficiency using an N– C swap design of donor and acceptor pairs, enabling comparison under conditions where the interfluorophore distance is effectively constrained. While only a single linker length was tested, this design minimizes distance-dependent variability and allows isolation of configurational effects. To ensure the robustness of the observed differences, FRET efficiency was evaluated using multiple complementary approaches, including acceptor photobleaching and sensitized emission measurements [9–11]. These methods yielded consistent results, confirming higher FRET efficiency in the AD configuration compared with the DA configuration. Furthermore, transformation of FRET efficiency into the relative parameter *K* = *E*/(1−*E*) [28,29] revealed that the ratio between configurations (orientation index: *OI* = *K*_AD_/*K*_DA_) was consistently and significantly greater than unity, supporting a systematic difference in energy transfer efficiency that is conserved across the measurement methods. Spectral overlap between donor and acceptor contributes to FRET efficiency [19,20,35]. EYFP, which exhibits a red-shifted absorption and emission tail compared with EGFP, possesses greater spectral overlap with mCherry [21,36], resulting in the higher FRET efficiency compared with EGFP in the present study. This observation confirms that, in addition to proximity, spectral properties are an important determinant of FRET beyond the subcellular compartment.

FRET efficiency is generally sensitive to local physicochemical conditions such as pH, ionic strength, temperature, and molecular crowding [20,21]. However, despite the distinct biochemical environments of the nucleus and cytoplasm [22,23], we observed comparable FRET efficiencies across these compartments for all constructs. Moreover, the superiority of the AD configuration over the DA configuration was consistently observed regardless of subcellular localization and donor spectral variants (EGFP and EYFP). These findings indicate that bulk cellular environmental factors are not the primary drivers of the differences observed in the present study.

Notably, the difference in FRET efficiency between DA and AD configurations in this study cannot be explained by donor-acceptor distance, static orientation, or cellular environment alone. The PAE and pLDDT analyses by AlphaFold2 suggested a substantial conformational flexibility of the fusion proteins, particularly in the linker regions, indicating that donor-acceptor geometry is dynamically sampled in living cells [37,38]. Such flexibility would lead to time-averaged distances and orientations rather than fixed values [3]. In this context, asymmetric conformational sampling between DA and AD configurations may result in differences in observed FRET efficiency in living cells. Furthermore, the limited rotational freedom of fluorescent proteins may bias dynamic orientation sampling, thereby contributing to orientation-dependent effects governed by N–C terminal topology [17,39].

In this study, AlphaFold2-based structural predictions showed highly conserved donor-acceptor distances across all models, ranging from 22.3 to 24.6 Å. These distances predicted near-saturated FRET efficiencies and therefore failed to discriminate those between DA and AD configurations. While orientation factors (*κ*²) were considered, calculations based on the AlphaFold models did not reproduce the experimentally observed differences in FRET efficiency. This discrepancy highlights the limitations of static structural models [40,41]. Consequently, our results suggest that FRET efficiency in live-cell systems cannot be fully explained by static structural parameters, but is likely influenced by dynamic conformational sampling and local interactions [42–44]. FRET efficiency (*E*) is determined by the competition between the energy transfer rates (*k_FRET_*) and the intrinsic decay rates of the donor, including both radiative (*k_f_*) and non-radiative (*k_nr_*) pathways: *E* = *k_FRET_*/(*k_FRET_* + *k_f_* + *k_nr_*) [3,6]. Given that donor fluorescence intensity showed no correlation with FRET efficiency in living cells in this study, variations in donor emission properties, including the radiative decay rate, are unlikely to be a primary determinant of the observed differences between DA and AD configurations. Taken together, our findings suggest that dynamic intramolecular factors, including distance fluctuations, orientation sampling, local photophysical environment, and transient interactions, collectively contribute to FRET efficiency in living cell systems (41, 42, 44). These factors may modulate both the rate of energy transfer and competing non-radiative decay pathways, thereby potentially influencing the observed FRET output in the present study.

To further investigate the potential basis of the experimentally observed DA/AD asymmetry in the FRET efficiencies, we performed ensemble FRET simulations incorporating conformational fluctuations and distance distributions. While the concept that conformational heterogeneity influences ensemble FRET measurements is well established [39,42–44], our approach differs in that experimentally measured FRET efficiencies in living cells were used to estimate the effective mean chromophore distances under defined conformational fluctuations, thereby enabling quantitative evaluation of DA/AD asymmetry. Our ensemble FRET simulations suggest that dynamic conformational sampling is a critical determinant of the effective mean chromophore distance required to reproduce experimentally observed FRET efficiencies. Notably, increasing conformational fluctuation (*σ*) progressively enhanced the predicted DA/AD distance asymmetry (Δ*μ*_eff_), suggesting that dynamic structural sampling amplifies the apparent asymmetry detected in live-cell FRET measurements. Thus, the *OI* provides an experimental measure of configurational asymmetry, while the ensemble FRET analysis provides a theoretical framework explaining how such asymmetry can emerge from conformational distance sampling. These findings suggest that the apparent DA/AD asymmetry in live-cell FRET efficiencies is influenced predominantly by dynamic conformational properties rather than by static conformational geometry alone [40,41].

While the present ensemble model provides a theoretical explanation for the experimentally observed asymmetry, several limitations should be acknowledged. In particular, the generality of the observed effect across different linker architectures remains to be established. The simulations assumed Gaussian distance distributions and did not explicitly describe the molecular motions responsible for these distributions. Furthermore, the AlphaFold models represent static conformations and therefore cannot capture the full range of structural dynamics occurring in living cells [40,41]. Future molecular dynamics simulations [38,42] combined with experimentally constrained structural analyses will be required to characterize the underlying conformational landscape in greater detail and determine how dynamic structural sampling contributes to FRET asymmetry. Additionally, while the present study used two independent intensity-based FRET approaches (acceptor photobleaching and sensitized emission measurements) that yielded consistent results, future studies with fluorescence lifetime-based FRET measurements would provide complementary validation using a methodology less dependent on fluorescence intensity measurements [45,46].

In summary, we demonstrated that the AD configuration consistently exhibited higher FRET efficiency than the DA configuration across donor spectral variants and cellular compartments. This difference cannot be explained by proximity or static structural models alone, but instead potentially reflects contributions from dynamic conformational behavior and local molecular interactions. The ensemble FRET simulations suggest that dynamic structural sampling amplifies the apparent DA/AD asymmetry observed in live-cell FRET measurements. Our findings highlight the importance of considering molecular fluctuations in live-cell environments and provide implications for the optimization of intramolecular FRET-based biosensors.

## Materials and Methods

### Plasmid construction

The pEGFP-C1 and pEYFP-C1 expression vectors were purchased from Clontech Takara Bio (Kusatsu, Japan). The pmCherry-NLS plasmid was a gift from Dr. Martin Offterdinger (Addgene plasmid #39319; URL: https://n2t.net/addgene:39319) [47].

To generate the pCMV-mCherry-MCS expression vector, which contains the same promoter, codon usage, and multiple cloning site (MCS) as the pEGFP-C1 and pEYFP-C1 vectors, we first deleted the sequence coding for *EYFP* (613–1329 bp) in pEYFP-C1 via inverse PCR using the KOD Plus mutagenesis kit (Toyobo, Osaka, Japan). This region was replaced with the *mCherry*-coding sequence amplified from the pmCherry-NLS vector using overlap extension PCR with KOD Plus Neo (Toyobo). To generate EGFP-mCherry and EYFP-mCherry expression vectors, the mCherry fragment harboring the GGSGGS linker at the 5′ end was amplified from pmCherry-NLS by PCR using KOD Plus Neo and inserted into the EcoRI and BamHI sites of pEGFP-C1 and pEYFP-C1. To generate mCherry-EGFP and mCherry-EYFP expression vectors, the EGFP or EYFP fragment together with the GGSGGS linker at the 5′ end was amplified from pEGFP-C1 or pEYFP-C1 by PCR using KOD Plus Neo and inserted into the EcoRI and BamHI sites in pCMV-mCherry-MCS.

The fluorescent proteins were fused with an identical 18-amino acid linker, and the stop codon of the N-terminal fluorophore and the start codon of the C-terminal fluorophore were deleted to generate continuous fusion proteins of 492 amino acids. The linker sequence, including the remaining MCS and the GGSGGS motif, was 5′-tcc gga ctc aga tct cga gct caa gct tcg aat tct ggt ggt tct ggt ggt tct-3′, which encodes the peptide sequence SGLRSRAQASNSGGSGGS. The resulting plasmids, EGFP-mCherry, EYFP-mCherry, mCherry-EGFP, and mCherry-EYFP, were verified by Sanger sequencing.

### Cell culture and transfection

COS-1 cells were originally obtained from ATCC (Manassas, VA, USA) and kindly provided by the laboratory at Department of Anatomy and Neurobiology, Graduate School of Medical Science, Kyoto Prefectural University of Medicine (Kyoto, Japan). Cells were cultured in Dulbecco’s Modified Eagle Medium (DMEM; Nacalai Tesque, Kyoto, Japan) supplemented with 5% fetal bovine serum (FBS; Biosera, Cholet, France) and 1× penicillin-streptomycin (Nacalai) under a 5% CO_2_/95% air atmosphere and 100% humidity at 37°C. All media in subsequent experiments contained 1× penicillin-streptomycin unless otherwise noted.

Prior to transfection, cells were seeded in 35-mm-diameter glass-bottom dishes (hole diameter: 15 mm, Violamo®, As One Corp., Osaka, Japan) pre-coated with 0.01% poly-L-lysine (Nacalai) at 2.5 × 10^4^/dish. The next day, the cells were transiently transfected with plasmids (500 ng/dish) using PEI-MAX^®^ (MW: 40,000, Ref # 24765-100, Polysciences Inc., Warrington, PA, USA) at 2 µg/dish diluted in Opti-MEM (Thermo Fisher Scientific, Waltham, MA, USA) for 3 h, followed by incubation in DMEM/5%FBS for recovery culture. For living cell imaging, the medium was replaced with phenol-red and bicarbonate free medium, Leibovitz’s L15 (Thermo Fisher Scientific) supplemented with 1 g/L D-glucose, and the cells were incubated at 37°C under normal air (CO_2_-free condition) and 100% humidity in the on-stage chamber (Tokai Hit, Shizuoka, Japan).

### Living cell FRET imaging

FRET microscopy was performed as described previously with minor modifications [8–10]. Fluorescence images of a single Z-section were acquired using a confocal laser-scanning microscope (FV3000, Olympus, Tokyo, Japan) at 1,024 × 1,024 pixels (scan speed: 8.0 µs/pixel) with 0.1% laser power at an excitation wavelength of 488 nm using a UPlanSApo 40×/0.95 NA objective lens (zoom factor: 4). During image acquisition, cells were maintained at 37°C in the on-stage incubation chamber. Donor fluorescence of EGFP or EYFP was detected at 500–540 nm upon 488-nm excitation (laser power: 0.1%, detector high voltage: 520, gain: 1,000); acceptor fluorescence of mCherry was detected at 570–620 nm (detector high voltage: 600, gain: 1,000). For acceptor photobleaching, the entire imaging field (1,024 × 1,024 pixels, zoom factor: 4) was irradiated with a 561-nm laser at 100% power (pixel dwell time: 2.0 µs) for 22.1 s to ensure that the entire cell was included within the bleached region. Scanning parameters and photobleaching settings were kept constant throughout all image acquisition procedures.

For the image data processing, OIR files were acquired using the FV3000 system and exported as 16-bit grayscale TIFF images in Raw Data Extracted mode with a fixed pixel dimension (1,024 × 1,024 pixels) to preserve original fluorescence intensity values. The converted TIFF images were analyzed using Zen lite software (version 3.13; Carl Zeiss, Jena, Germany). Regions of interest (ROIs) were defined as circular areas with a 50-pixel diameter and placed separately within the nucleus or cytoplasm (up to three ROIs per nucleus or cytoplasm each). Fluorescence intensities from the ROIs were averaged to determine representative values for the nuclear and cytoplasmic compartments of each cell. The overall mean fluorescence intensity per cell was calculated by averaging the nuclear and cytoplasmic values.

### Evaluation of FRET efficiencies

The FRET efficiency determined by acceptor photobleaching (*E_pb_*) was calculated using the following equation:

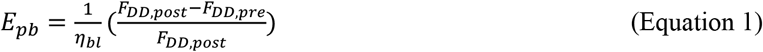

where *F_DD,pre_* and *F_DD,post_* represent the donor fluorescence intensities under donor excitation before and after acceptor photobleaching, respectively, and *η_bl_* represents the bleaching efficiency, defined as the fraction of the acceptor fluorophore that was photobleached (Supplementary Fig. S1).

The FRET efficiency for sensitized emission measurements (*E_se_*) was calculated as follows.

First, the fluorescence ratio was defined as:

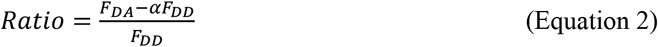

where *F_DA_* and *F_DD_* represent the fluorescence intensities in the acceptor and donor channels under donor excitation, respectively, and *α* denotes the donor bleed-through coefficient [4,48]. Since direct acceptor excitation was not detected in this system, no correction for acceptor bleed-through was applied.

The *E_se_* was then calculated as:

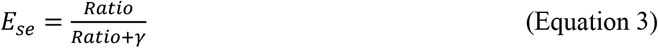

where γ is a correction factor defined as previously described [48,49]:

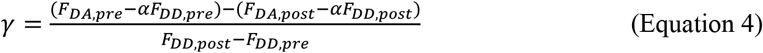

fluorescence intensities measured before and after acceptor photobleaching, respectively.

All fluorescence intensities, bleed-through coefficients, and bleaching efficiencies were corrected by subtracting background fluorescence from a cell-free region prior to calculation.

### Analyses of 3D protein structure models

The 3D structures of the four fusion proteins (EGFP-mCherry, mCherry-EGFP, EYFP-mCherry, and mCherry-EYFP) were predicted using AlphaFold2 (DeepMind Technologies, London, UK) [24], implemented via ColabFold (Steinegger Lab, Seoul National University, Seoul, South Korea) [32]. The predicted structures were visualized and analyzed using UCSF ChimeraX (URL: https://www.cgl.ucsf.edu/chimerax/; version 1.11, University of California, San Francisco, CA, USA) [33]. For each construct, five structural models were generated and ranked based on the pLDDT scores. In this study, the top-ranked model was used for structural analysis, except for inter-chromophore distances, which were averaged across all five models (Supplementary Fig. S4a, b).

The chromophore was approximated using the aromatic ring of the tyrosine residue that constitutes its core (Tyr67 and Tyr328 in the DA constructs; Tyr72 and Tyr320 in the AD constructs; amino acid numbering refers to the full-length fusion protein of 492 amino acids). Distance between chromophores (*R*) was defined as the distance between the Cζ atoms of the tyrosine residues [12].

The interplanar angles were calculated using planes defined by the Cγ, Cδ1, and Cδ2 atoms of the tyrosine aromatic rings.

The orientation factor (*κ*²) was estimated based on the relative orientation of the transition dipole moments of the chromophores, with dipole vectors approximated using Cγ-Cζ vectors of the tyrosine aromatic rings. *κ*² was calculated using the standard geometric definition as follows:

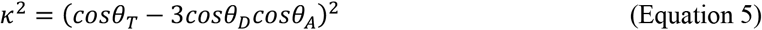

where *θ_T_* represents the angle between the transition dipole moments of the donor and acceptor derived from the chromophore geometry, and *θ_D_* and *θ_A_* are the angles between the respective dipole moments and the interchromophore vector [3,17].

Theoretical FRET efficiencies (*E_th_*) were calculated using the Förster equation [35]:

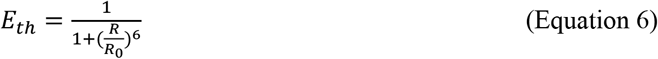

where *R* represents the donor-acceptor distance measured between Cζ atoms of tyrosine residues constituting chromophore cores in ChimeraX. The Förster radius (*R*_0_) was corrected using the estimated orientation factor (*κ*²) according to:

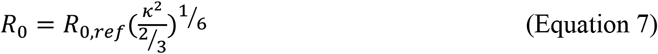

where *R*_0,ref_ is the reference Förster radius assuming *κ*^2^ = 2/3 obtained using the FPbase FRET Calculator (accessed May 9, 2026, https://www.fpbase.org/fret/) [36].

### Ensemble FRET simulation

To investigate the influence of conformational fluctuations on the experimentally observed FRET efficiencies, an ensemble FRET model was implemented in Wolfram Mathematica 15.0 (Wolfram Research, Champaign, IL, USA) (Supplementary Code S1) [34]. The chromophore distance was modeled as a Gaussian distribution [39]:

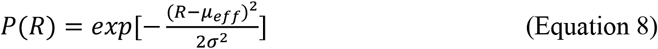

where *μ*_eff_ is the effective mean chromophore distance and *σ* represents the assumed extent of conformational fluctuation [44]. For each construct, the effective mean chromophore distance (*μ*_eff_) was defined as the theoretical mean chromophore distance required to reproduce the experimentally observed FRET efficiency for a given *σ*.

Because the ensemble FRET efficiency was calculated as the ratio of two integrals, the normalization constant of the Gaussian distribution cancels and was therefore omitted. Numerical integration was performed over a chromophore distance range of 0–150 Å. This integration range was sufficient to ensure numerical convergence for the conformational fluctuations analyzed in this study (*σ* ≤ 50 Å).

The ensemble FRET efficiency was calculated as the distance-distribution-weighted average [39,42,43],

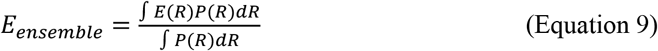

where the distance-dependent FRET efficiency is given by

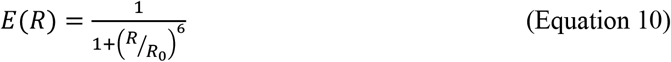

The value of *μ*_eff_ was determined numerically using the FindRoot function implemented in Wolfram Mathematica.

Contour maps representing ensemble FRET efficiencies were generated for each construct. The contour corresponding to the experimentally observed FRET efficiency was subsequently used to determine the effective mean chromophore distance (*μ*_eff_) for a given value of *σ*.

The effective distance asymmetry was defined as

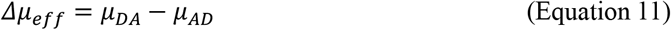

The dependence of Δ*μ*_eff_ on the assumed conformational fluctuation (*σ*) was subsequently evaluated.

## Statistical analysis

Data in Figs. 2–4 are expressed as mean ± SE unless otherwise indicated. These data were analyzed by a one-sample *t*-test, unpaired *t*-test, or one-way ANOVA for inter-chromophore distance and two-way ANOVA for FRET efficiency, both followed by Bonferroni/Dunn’s post-*hoc* test. Statistical analyses were performed using StatView version 5.0 (SAS Institute Inc., Cary, NC, USA). A *p* value < 0.05 was considered statistically significant.

For *t*-tests, the sign of the *t*-statistic reflects the order in which the two groups were compared. *F*-statistics are reported as *F*(df1, df2), where df1 and df2 indicate the numerator and denominator degrees of freedom, respectively. The *t*-statistics are reported as *t*(df), where df indicates the degrees of freedom.

Linear regression analysis was performed using Microsoft Excel, and coefficients of determination (*r*²) were obtained from the fitted trendlines.

## Code availability

The custom Mathematica scripts implementing the ensemble FRET simulations described in this study are provided as Supplementary Code S1 and are also available on Zenodo (URL: https://doi.org/10.5281/zenodo.21991881) [34]. All distance-related calculations in the Mathematica notebook were performed in nm, whereas distances are reported in Å in the manuscript and Figures.

## Acknowledgements

We thank Dr. Martin Offterdinger for providing the pmCherry-NLS plasmid from Addgene (#39319). COS-1 cells were obtained from stocks maintained in the Department of Anatomy and Neurobiology, Graduate School of Medical Science, Kyoto Prefectural University of Medicine. We also thank Gabrielle White Wolf, Ph.D., from Edanz (https://jp.edanz.com/ac) for editing a draft of this manuscript.

## Author contributions

T.T. conceived the idea and designed the study. T.T. performed all experiments, data collection, structural bioinformatic analysis, and ensemble FRET simulations. T.T. and M.R.G. conducted statistical evaluations. T.T., M.R.G., and T.N. interpreted the data. T.T. drafted the manuscript with input from M.R.G. and T.N. All authors approved the final version of the manuscript.

## Data availability statement

All relevant data are included in this article. Any additional data associated with the experiments are available from the corresponding author upon reasonable request.

## Supplementary Figure Legends

**Figure S1.** Bleaching efficiency for acceptor photobleaching FRET microscopy in living cells. a. Confocal live-cell images of COS-1 cells expressing mCherry before (*Pre*) and after (*Post*) bleaching using 561-nm laser with a maximum power. Raw and pseudocolored images are provided in the *top* and *bottom,* respectively. Pseudocolor bars indicate fluorescence intensity, with H and L representing high and low signal intensity, respectively. Bar = 20 µm. b. Relative fluorescence intensity of ROIs placed in the nucleus and cytoplasm of COS-1 cells expressing mCherry before (*Pre*) and after (*Post*) bleaching with 561-nm laser. Values are expressed as mean ± SE (n = 16 for nucleus; n = 14 for cytoplasm). c. Bleaching efficiency within the nucleus (*Nuc*) and cytoplasm (*Cyt*) calculated from the fluorescence intensity of COS-1 cells expressing mCherry before and after beaching with 561-nm laser with maximum power. Values are expressed as mean ± SE (n = 16 for nucleus; n = 14 for cytoplasm).

**Figure S2.** Sequence coverage maps for EGFP-mCherry (a), mCherry-EGFP (b), EYFP-mCherry (c), and mCherry-EYFP (d) generated by AlphaFold2. The horizontal axis represents amino acid positions, and the vertical axis represents sequences in the multiple sequence alignment (MSA). Colors indicate sequence identity to the query sequence, while the black line represents MSA coverage across residue positions. The artificial linker region (approximately residues 230–260) shows markedly lower sequence coverage than the fluorescent protein domains, reflecting the limited availability of homologous sequences.

**Figure S3.** Predicted aligned error (PAE) plots for the top five ranked structures of EGFP-mCherry (a), mCherry-EGFP (b), EYFP-mCherry (c), and mCherry-EYFP (d) generated by AlphaFold2. The PAE plots indicate the predicted accuracy of relative positions between residue pairs, with lower PAE values (blue) representing higher confidence. Consistently low PAE values within the fluorescent protein domains across the top five ranked models indicate high confidence in the predicted domain structures, whereas relatively higher PAE values between the two domains reflect greater uncertainty in their relative positioning.

**Figure S4.** Theoretical structural metrics and FRET efficiencies derived from ColabFold and UCSF ChimeraX analyses. a. Inter-chromophore distances measured between Tyr-Tyr Cζ atoms for each construct, shown as the average of the top five ranked AlphaFold2 models. Data are presented as mean ± SE (n = 5 per construct). One-way ANOVA followed by Bonferroni/Dunn’s post*-hoc* test. n.s., not significant. b. Tabulated inter-chromophore distances (Å) for the top five ranked structures of each construct. c. Theoretical FRET efficiencies calculated for each construct across a range of orientation factors (*κ*^2^) from 0 to 4, including the isotropic assumption value (*κ*² = 2/3). *κ*² = 0 was treated as not applicable (n.a.) because energy transfer is theoretically prohibited under orthogonal dipole orientation. The dashed line indicates 100% FRET efficiency.

