## Supplemental Figures for "Topology-dependent FRET efficiency in living cells via N-C swapping of fluorescent protein fusions" (from another version of this preprint: version 1, 2026-06-06)

Supplementary Figure S1

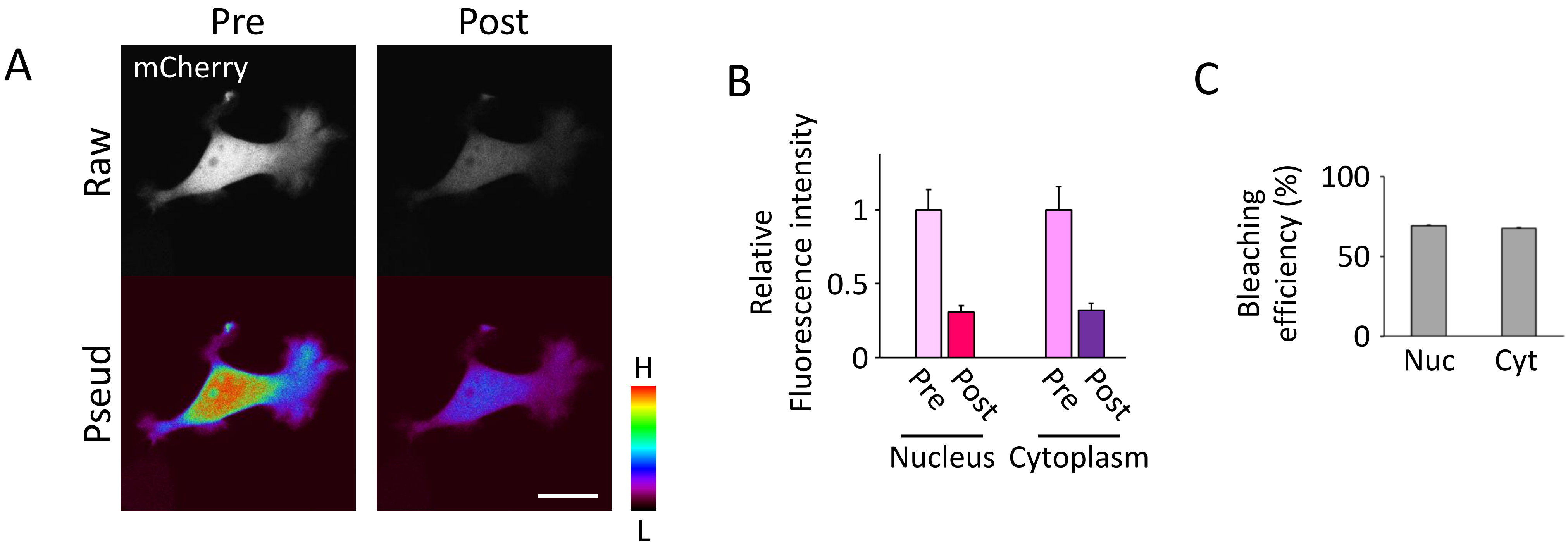

Supplementary Figure S2

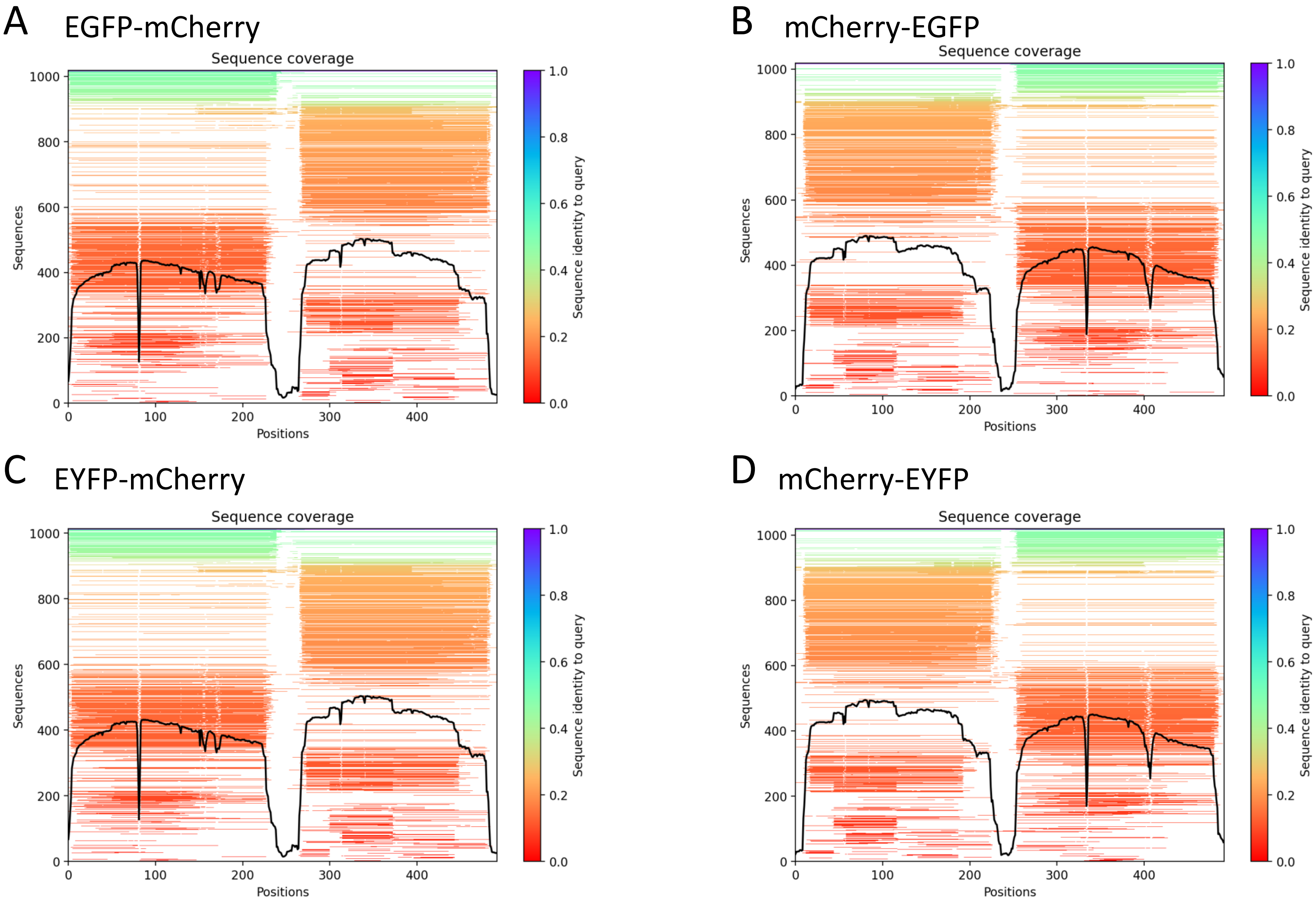

Supplementary Figure S3

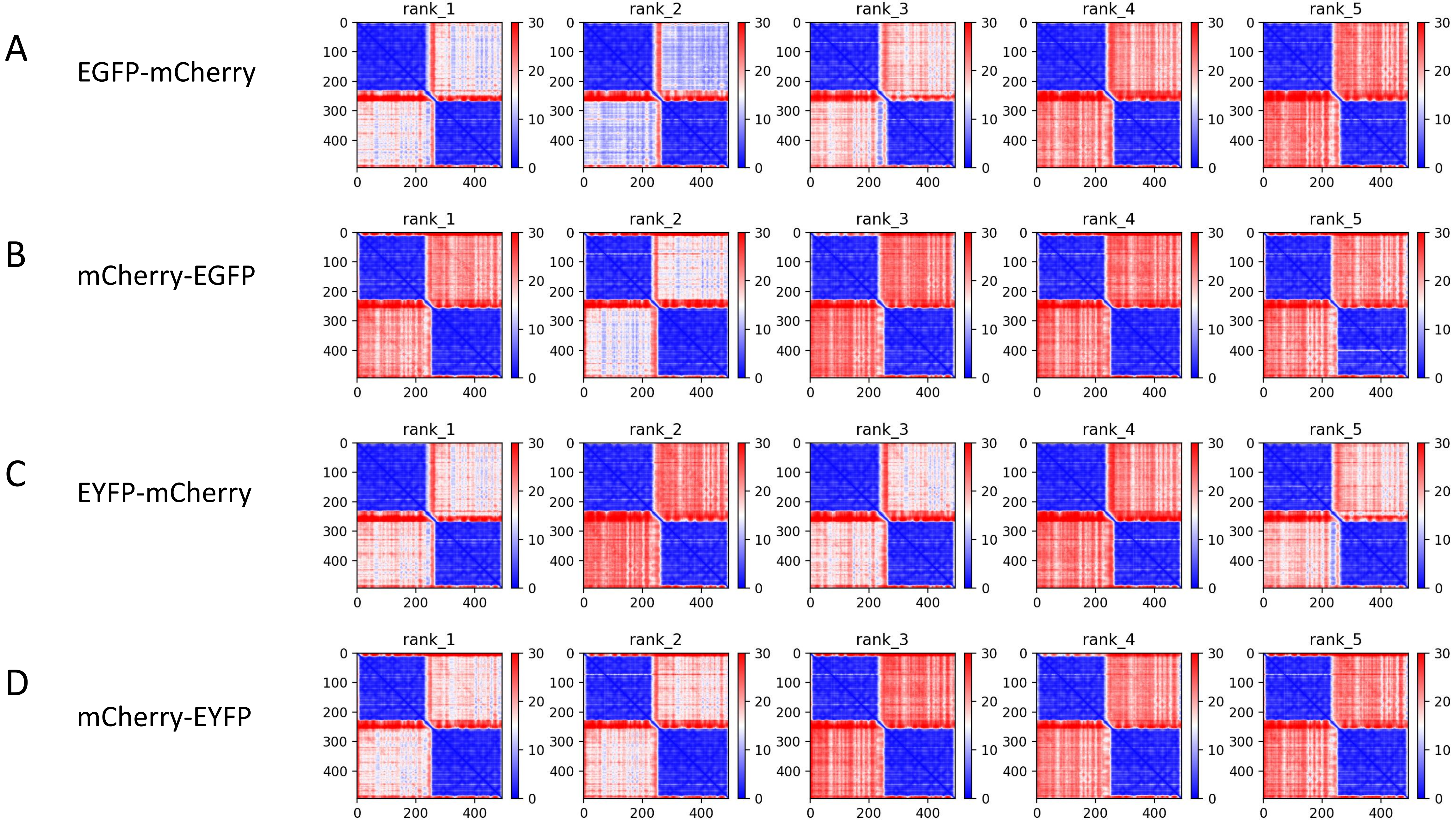

Supplementary Figure S4

A

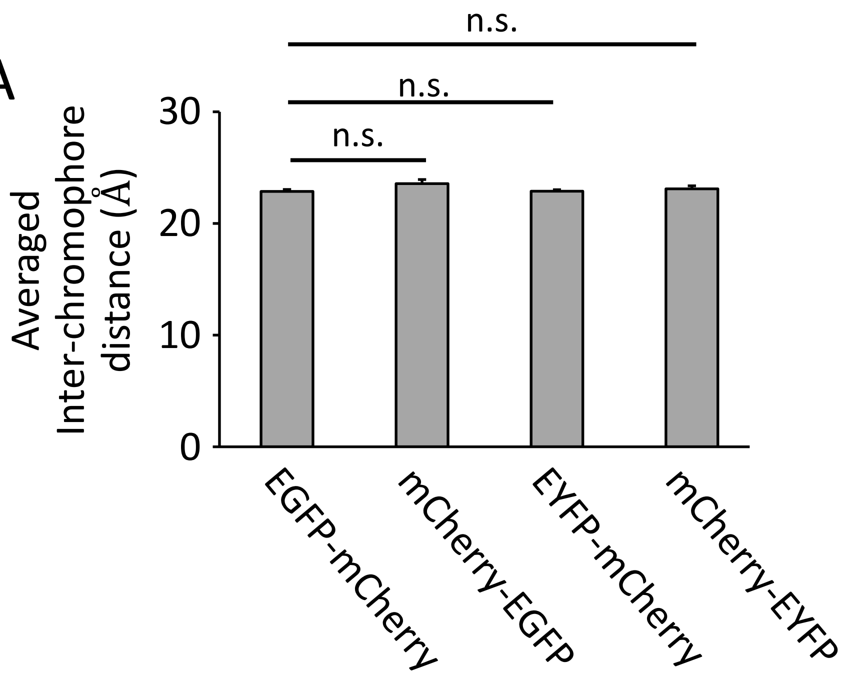


B

Averaged inter-chromophore distances (Å) in the five top-ranked structures

| | EGFP-mCherry | mCherry-EGFP | EYFP-mCherry | mCherry-EYFP |
|---|---|---|---|---|
| rank_001 | 22.9 | 24.1 | 22.8 | 23.4 |
| rank_002 | 22.9 | 23.6 | 22.9 | 23.2 |
| rank_003 | 23.5 | 24.6 | 23.0 | 22.9 |
| rank_004 | 22.5 | 23.1 | 22.5 | 23.9 |
| rank_005 | 22.6 | 22.4 | 23.3 | 22.3 |

C

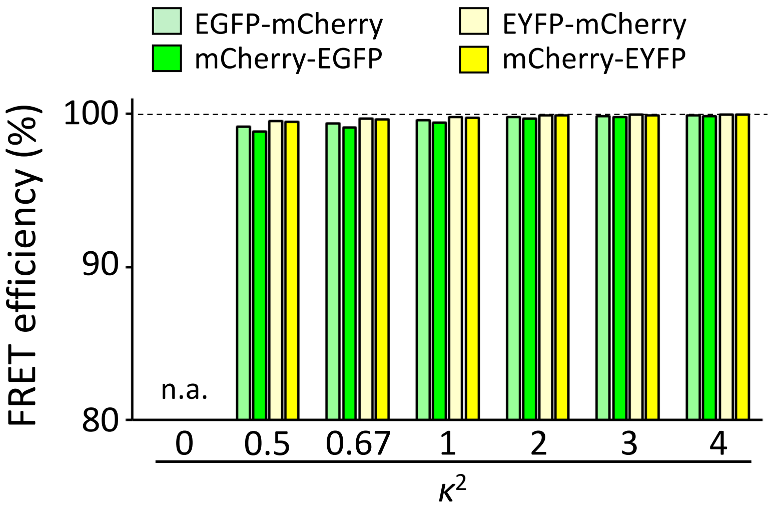
